# The NADPH oxidase and microbial killing by neutrophils, with a particular emphasis on the proposed antimicrobial role of myeloperoxidase within the phagocytic vacuole

**DOI:** 10.1101/019174

**Authors:** Adam P. Levine, Anthony W. Segal

## Abstract

This review is devoted to a consideration of the way in which the NADPH oxidase of neutrophils, NOX2, functions to enable the efficient killing of bacteria and fungi. It includes a critical examination of the current dogma that its primary purpose is the generation of hydrogen peroxide as substrate for myeloperoxide catalysed generation of hypochlorite. Instead it is demonstrated that NADPH oxidase functions to optimise the ionic and pH conditions within the vacuole for the solubilisation and optimal activity of the proteins released into this compartment from the cytoplasmic granules, which kill and digest the microbes. The general role of other NOX systems as electrochemical generators to alter the pH and ionic composition in compartments on either side of a membrane in plants and animals will also be examined.

## 1 The respiratory burst of professional phagocytic cells is accomplished through the action of the NADPH oxidase, NOX2

In 1933 Baldridge and Gerard observed the ‘extra respiration of phagocytosis’ when dog leukocytes were mixed with Gram positive bacteria, and assumed that it was associated with the production of energy required for engulfment of the organisms. It was later shown that this ‘respiratory burst’ was not inhibited by the mitochondrial poisons, cyanide [1] or azide [2], which indicated this is a non-mitochondrial process. The hunt for the neutrophil oxidase was then on because oxidative phosphorylation is by far and away the major mechanism by which oxygen is consumed in mammalian cells, and another system that could consume oxygen at a similar rate was of considerable interest. Although many oxidases use NADH or NADPH as substrate, this oxidase was specifically called the NADPH oxidase because in the early days there was quite a controversy as to whether the substrate was NADH [3] or NADPH [4] and the matter was finally settled in favour of NADPH.

In 1957 a new disease called ‘fatal granulomatosus of childhood’ was discovered [5], [6]. Neutrophils from subjects with this ‘chronic granulomatous disease’, or CGD, demonstrated an impaired ability to kill *Staphylococcus aureus*, as well as an absent respiratory burst, thereby linking the NADPH oxidase to the bacterial killing process.

The next big development was the discovery of an enzyme, superoxide dismutase (SOD), which rapidly converted superoxide O_2_^-^ to H_2_O_2_ [7]. The inclusion of SOD allowed the detection of superoxide (O_2_^-^) in biological systems because it specifically inhibited superoxide induced reactions. The use of this enzyme enabled Babior and colleagues to demonstrate that activated neutrophils generate O_2_^-^, which was in agreement with the previous observation that these cells generate hydrogen peroxide (H_2_O_2_) [8] which is produced by the dismutation of O_2_^-^.

The search was then on for an ‘enzyme’ that would transport an electron from NADPH to oxygen to form O_2_^-^. This turned out to be a demanding task because the activity disappeared with simple purification techniques, for reasons that will become obvious later. In 1978 we discovered a very low potential cytochrome b that was absent from neutrophils of patients with CGD, which appeared to fulfil the requirements of the oxidase [9], [10]. Borregaard and colleagues [11] found the cytochrome b in patients with the autosomal pattern of inheritance, and surprisingly also in another such patient with the X-linked pattern of inheritance, which led them to question the relationship between the cytochrome b and the oxidase. We went on to confirm that patients with autosomal recessive CGD did in fact have the cytochrome b in their neutrophils, but were unable to transfer electrons on to it when the cells were activated [12], indicating the absence of an upstream electron transporting molecule, or of an activation mechanism. The latter case turned out to be the correct one. Over time it was demonstrated that the cytochrome comprised gp91^phox^, the electron transporting component, together with another membrane component, p22^phox^ [13], [14] in stoichiometric equivalence [15]. In addition, five cytosolic proteins were required, p47^phox^ (also called NOX organiser 1 [16]), p67^phox^ (NOX activator 1 [16]) [17], [18], p40^phox^ [19] and the small GTP binding protein Rac which dissociates from its binding partner GDI [20]. All of these components, apart from GDI, move to the membranes and activate electron transport through the cytochrome. The requirement for the co-ordinated integration of these multiple components for an active oxidase system explains why it proved impossible to obtain the active oxidase by classical biochemical purification techniques, because these resulted in dissociation of the complex with loss of enzymic activity.

It proved possible to recombine most of the components of the oxidase in a ‘cell free’ assay [21], [22] in which solubilised membranes, or purified cytochrome b, could be mixed with cytosol, or the purified cytosolic protein components, and NADPH. Electron transport was then induced by the addition of arachidonic acid, or a detergent, which must have changed the conformation of the cytochrome, giving it access to substrate and the accessory binding proteins. This ‘cell free’ assay system was also useful in characterising some cases of autosomal recessive CGD, where the missing, or abnormal, cytosolic components of the oxidase were identified by complementation of the deficient cytosol with semi-pure proteins [23].

Once the cytochrome b had been identified, it could be isolated by following its spectro-graphic signature [24] through the different stages of purification. We showed that it was a flavocytochrome, and characterised it in terms of its NADPH, FAD [25], haem [26] and carbohydrate [27] binding sites. We purified, and obtained amino acid sequence from, gp91^phox^ [28] and demonstrated that it was coded for by the gene, abnormalities of which cause X-CGD, that had been cloned based on its chromosomal localisation [29].

## 2 The relationship between the phagocyte oxidase and microbial killing

It was known that stimulated neutrophils generated H_2_O_2_ and that they contained a very high concentration of a peroxidase, myeloperoxidase (MPO) [30], in their granules [31]. Klebanoff demonstrated the fixation of iodine by neutrophils phagocytosing bacteria, and that concentrated MPO can kill *E. coli* in combination with iodide and H_2_O_2_ *in vitro*, suggesting that this may contribute to bacterial killing by neutrophils *in vivo* [32]. The discovery that absence of the oxidase in CGD was accompanied by greatly increased incidence of clinical infections, together with a clear impairment of bacterial killing by neutrophils from these patients, established the requirement of the activity of this oxidase for efficient microbial killing. Defective iodination by their neutrophils reinforced the feasibility of a physiological role for the MPO-H_2_O_2_-iodide antimicrobial system [33].

The demonstration that activated neutrophils can generate O_2_^-^ [34] produced enormous excitement. Superoxide, H_2_O_2_ and their reaction products were proposed as effector molecules that could themselves cause the microbicidal lethal reactions. In addition, because of the predicted toxicity of these molecules [35] that could be generated by neutrophils, and because neutrophils were found in abundance in inflammatory sites where tissue damage occurred, it was a natural progression to suggest that the tissue damage was being produced by the neutrophil generated radicals [36]. This implied that antioxidant molecules that reacted with, and consumed, these reactive oxygen molecules might provide valuable therapeutic anti-inflammatory agents [37].

## 3 The case for and against a primary role for myeloperoxidase in the killing of ingested microbes within the phagocytic vacuole

### 3.1 The myeloperoxidase-halide-hydrogen peroxide antibacterial system

Much evidence has been generated to support the concept that MPO oxidises chloride and other halides to generate their hypohalous acids, and that these compounds kill the organisms within the phagocytic vacuole. It is important however to be aware of two important factors. The first is that these theories originated when it was believed that the pH within the phagocytic vacuole of the neutrophil is acidic, with a pH of 6.5, falling to 4.0 [38]. The *in vitro* MPO-halide-hydrogen peroxide antibacterial system is critically dependent upon the ambient pH, being highly efficient at pH 5.0 and ineffective at pHs of 7.0 and above [32], [39]. Newly obtained data demonstrate that in normal human neutrophils the respiratory burst elevates the pH within the phagocytic vacuole to *≈*9.0, at which pH the *in vitro* peroxidatic effect of MPO is almost non-existent [40]. The second issue of consequence is that in the test tube it is exceedingly difficult to accurately distinguish between organisms that have been fully engulfed into a closed vacuole, and those that remain in suspension in the medium or attached to the neutrophils but not fully taken up. The latter situation is particularly relevant when dealing with bacteria that have a tendency to form microcolonies and biofilms, such as *Staphylococci* [41], *Burkholderia cenocepacia* and *Pseudomonas* [42], and *Serratia* [43], and fungi like *Aspergillus* [44] and *Candida* [45].

### 3.2 The consequences of generating H_2_O_2_ in the phagocytic vacuole

The NADPH oxidase generates O_2_^-^ in the vacuole, which naturally dismutates to H_2_O_2_ and this process is accelerated by MPO, which has superoxide dismutase activity [40], [46]. There is very strong evidence for the generation of H_2_O_2_ in the vacuole [47] as initially proposed on the basis of the oxidation of formate to CO_2_ [8]. It is not clear what then happens to the H_2_O_2_. The dogma is that it is used as substrate for MPO to generate HOCl, but this is very unlikely at the pH of 9.0 recently demonstrated in the neutrophil vacuole, at which the peroxidatic and chlorinating activities of MPO were shown to be very low [40]. An alternative scenario is that MPO acts first as a superoxide dismutase and then as a catalase [40], [48] to safely remove the H_2_O_2_ and regenerate its SOD activity, thereby preventing the damaging oxidation of the important granule proteins. It seems highly improbable that the neutrophil would synthesise, store and release highly specialised enzymes into the phagocytic vacuole, only to have them degraded by HOCl [49], [50]. One piece of evidence used to support the MPO-halide-hydrogen peroxide antibacterial system has been the misconception that patients with CGD are less susceptible to infections with catalase negative than positive organisms, because it was supposed that the former produced H_2_O_2_ that was then utilised by the MPO system for their own destruction. This was in fact shown not to be the case as catalase negative *S. aureus* [51] and *Aspergillus nidulans* [52] were found to be just as virulent as the catalase positive strains in CGD mouse models.

### 3.3 Evidence for the generation of HOCl in the phagocytic vacuole

Several attempts have been made to measure HOCl in the vacuole [53]. Hurst’s group used fluorescein coupled to polyacrylamide microspheres to generate fluorescent reporter groups that could then be uncoupled from the particles by reduction of the cystamide disulphide bond. The mono-, di-and trichlorofluorescein compounds were separated and measured by liquid chromatography. They demonstrated the generation of these chlorinated species by the MPO-H_2_O_2_-Cl^-^ system had a pH optimum of 6-7.5 with virtually no activity at physiological pH of *>*8.5 [54]. They assessed the efficiency of conversion of O_2_ to HOCl at about 11%, which they judged to all be within closed vacuoles because the fluorescence was not quenched by extracellular methyl viologen, which effectively quenched the fluorescence of the beads alone. However, some of these quenching experiments were performed in 100% serum which we have found to largely abolish the quenching of the fluorescence of FITC labelled bacteria by methyl viologen (unpublished).

Various other attempts have been made to develop HOCl specific probes, for example that by Koide and colleagues [55], but once again specificity was not demonstrated under vacuolar conditions. They demonstrated fluorescence of their intravacuolar probe, but this did not occur by 90 seconds after phagocytosis of opsonised zymosan, by which time the respiratory burst would have been well advanced if not completed [40].

Painter and colleagues metabolically labelled *Pseudomonas* with [13C9]-L-tyrosine which were opsonised with serum and mixed with neutrophils in the presence of catalase and taurine, which were included to suppress extracellular, but not intracellular, chlorination. The, largely, ingested bacteria, were subjected to protease hydrolysis and levels of 13C9-derived chlorotyrosine and tyrosine were determined by gas chromatography and mass spectrometry (GC-MS) using the isotope dilution technique. Because neutrophil-derived tyrosine contains no [13C9]-L-tyrosine or its derivatives, this is a direct measure of the chlorination of intra-cellular *Pseudomonas*-derived tyrosine. They did find these products, but only a very small amount, about 0.15%, of the [13C9]-L-tyrosine was chlorinated [56], [57].

Chapman and colleagues [58] also used GC-MS to measure chlorinated tyrosine compounds. They found that about 0.2% of the tyrosines became chlorinated and that 94% of chlorination was of neutrophil rather than bacterial tyrosine residues even though tyrosine residues in bacteria were chlorinated approximately 2.5 times as efficiently as those in neutrophils. An interesting, and unexplained, observation in this study was that when azide was added as an inhibitor of MPO, it had exactly the opposite effect to that expected, resulting in a threefold increase in chlorination (their Figure 8). This group subsequently performed similar experiments with purified phagocytic vacuoles containing magnetic beads and achieved a chlorination efficiency of 1.1% of the vacuolar tyrosines [59].

We examined the targets of iodination after bacteria were phagocytosed [60]. Less than 0.3% of the oxygen consumed was utilised for iodination and almost all the iodine was incorporated into neutrophil proteins rather than those of the engulfed bacteria [61].

So how can we explain these results where some evidence of chlorination is observed while at the same time we know that conditions within the vacuole are most unfavourable for such chlorination reactions? We know that under certain experimental conditions up to one half of all phagocytic vacuoles can fail to close [62], [63] and in these vacuoles that communicate with the extracellular medium the MPO would be interacting with H_2_O_2_ and Cl^-^ at the pH of this extracellular medium, which is at about 7.4. The failure of closure of only a small proportion of the vacuoles could give rise to the observed chlorination results. It is also possible, as will be discussed below, that MPO has a dual function. In the closed vacuole it may act as a catalase and superoxide dismutase and that when the vacuole is unable to close it acts as a peroxidase.

### 3.4 Superoxide, H_2_O_2_ and HOCl are not as microbicidal as previously thought

The discovery of the production of O_2_^-^ by neutrophils was associated with the expectation that this free radical might itself be microbicidal [34], but it is now generally accepted that neither O_2_^-^ nor its dismutation product, H_2_O_2_, are damaging enough to be directly microbicidal [64]-[66], and as a result the emphasis has been placed upon the toxicity of HOCl generated by MPO.

We conducted experiments to directly determine the antimicrobial actions of O_2_^-^, H_2_O_2_ and HOCl on *S. aureus* and *E. coli* [61] on their own, and in the presence of granule proteins. It is important to assess their effects in the presence of the granule proteins because these are the circumstances under which these products of the oxidase would be generated. HOCl would not be produced in the absence of MPO, indicating that degranulation would have occurred into the vacuole where the bulk of the chlorination is observed [59] and where the granule proteins have been shown to be chlorinated [58].

In the presence of 25 mg/ml of granule proteins, which, whilst being a practical experimental concentration is almost certainly an underestimate of that pertaining in the vacuole, neither 100 mM O_2_^-^ or H_2_O_2_, nor 1mM HOCl killed either organism at neutral pH.

### 3.5 Myeloperoxidase deficiency

If the main function of the oxidase is to produce H_2_O_2_ as substrate for MPO mediated generation of HOCl, and if HOCl is the major microbicidal species, then MPO deficiency would be expected to predispose to serious infection. This does not turn out to be the case. In large studies involving between 57,000 and 70,000 individuals in America, Japan and Germany the incidence of MPO deficiency was found to be between 0.05 and 0.15% [67]-[69]. The incidence of infection in these subjects was very low. In the largest, German, study the distribution of diseases in the deficient patients did not differ from that of the general hospital population and only about 6% of these subjects had infectious disease. The MPO knock-out mouse demonstrated variable sensitivity to infection with a range of different organisms [70]. MPO-deficient and control mice were infected intranasally with various fungi and bacteria, and the number of residual microorganisms in the lungs was compared 48 hours later. MPO-deficient mice showed severely reduced cytotoxicity to *Candida albicans*, *Candida tropicalis*, *Trichosporon asahii*, and *Pseudomonas aeruginosa*. However, the mutant mice showed a slight but significantly delayed clearance of *Aspergillus fumigatus* and *Klebsiella pneumoniae* and had comparable levels of resistance to the wild type against *Candida glabrata*, *Cryptococcus neoformans*, *Staphylococcus aureus*, and *Streptococcus pneumoniae*. It is of note that infection with these agents generally occurs by multiplication from a relatively small inoculum rather than the instillation in massive numbers as a bolus, and that those organisms that were eliminated more slowly from the MPO deficient lungs tend to grow in clumps or mycelia that would be difficult for neutrophils to fully engulf (*vide infra*).

Whereas few, if any, MPO deficient subjects develop serious pyogenic bacterial or fungal infections, the penetrance of the molecular lesions of CGD is almost complete. Even though the rare individual with CGD has been reported as presenting in later life [71], *>*90% of patients present by the age of 20 years [72], and we are unaware of an individual with the molecular lesion of CGD, i.e. the relative of an affected patient, that did not develop clinically serious infections. The knock-out mice mirror the discrepancy observed in humans. CGD mice demonstrate a major vulnerability to infections with a wide variety of organisms whilst the propensity is only slightly increased in the MPO deficient mouse [73], [74]. This argues that the function of the NADPH oxidase cannot simply be to generate H_2_O_2_ as substrate for MPO dependent generation of HOCl. The MPO deficient mice appear to have a particular problem resisting infection with organisms like *Candida*, *Aspergillus* and *Pseudomonas* that form clumps, hyphae or biofilms. It is possible that these organisms are not phagocytosed and that they produce a state of frustrated phagocytosis in which the MPO acts at the pH of the extracellular medium, which is acid at sites of infection [75], under which circumstances HOCl could be produced and the extracellular organisms killed.

Approaching the issue from the aspect of a diagnostic laboratory investigating leukocyte function in patients with pyogenic infections [76], of 81 patients with a diagnosis of a primary phagocytic disorder, 48 had CGD and only two had MPO deficiency, of whom one had other neutrophil abnormalities as evidenced by delayed separation of the umbilical cord and severely impaired chemotaxis, and the other the relatively mild infection of furunculosis. The incidence of CGD is at most about 1 in 250,000 and that of MPO deficiency about 125 times as common, thus the predisposition to pyogenic infection in CGD is about 6,000 times as great as that of MPO deficiency. It thus seems totally implausible that MPO acts as the effector system for H_2_O_2_ produced by the NADPH oxidase that is defective in CGD.

## 4 Granule proteins

The unique structural characteristic of neutrophils is their content of cytoplasmic granules that comprise about 10% of the total cellular protein. The antimicrobial proteins and peptides are largely contained within the azurophilic granules where they are almost all strongly cationic, and are bound to a negatively charged sulphated proteoglycan matrix [77]. The granule contents comprise a diverse complement of enzymes and peptides, that have evolved to kill a wide variety of microorganisms and to digest diverse organic material [78], [79]. The most abundant of these enzymes are the neutral proteases, cathepsin G, elastase and proteinase 3, lysozyme, MPO and the defensins, but there are a host of other enzymes and proteins [80].

In the absence of clear evidence of microbial killing by products of the oxidase we investigated the possibility that the neutral proteases might be involved [81], which was particularly pertinent in view of the observation that we had made that the phagocytic vacuole was alkalinised by the oxidase [82] (as will be discussed later). We knocked out the genes for cathepsin G, and for elastase, and examined the susceptibility of the single and double knock-out mice to infection, and the competence of their neutrophils to kill bacteria and fungi *in vitro*. Despite normal neutrophil development and recruitment, the mice lacking either of these proteins were more susceptible to infection with *Aspergillus fumigatus*, and they were even more susceptible when deficient in both enzymes [83]. Differential responses were seen when these animals were challenged with *S. aureus* or *C. albicans* [84]. The elastase deficient mice were normally resistant to *S.aureus* but unduly sensitive to *C. albicans*, and the opposite was true for cathepsin G deficiency. The double knock-out mice were susceptible to both organisms. *In vitro*, the purified neutrophils from susceptible animals exhibited deficient microbial killing. These findings provided a clear example of selectivity of the microbicidal activity of the granule enzymes for different organisms. The same mice were used by others to demonstrate the requirement for cathepsin G, and to a greater extent both cathepsin G and elastase, for resistance to infection with *Streptococcus pneumonia* [85] or with mycobacteria [86].

The susceptibility of the mice made deficient in the neutral proteases was observed despite the presence of a normal respiratory burst, and of iodination, further evidence against a microbicidal action of the products of the oxidase or MPO [84].

Further evidence for the role of the neutral proteases was that selective inhibitors impaired the killing of *S. pneumonia* by human neutrophils, and purified elastase and cathepsin G were independently able to kill these organisms in *in vitro* assays [87].

Another set of protease deficient mice were made. These elastase deficient mice were found to be abnormally vulnerable to infection with *Klebsiella pneumoniae* and *E. coli* but resistant to *S. aureus* [88] whilst the cathepsin G deficient mice were resistant to all three organisms [89]. Mice lacking both cathepsin G and elastase demonstrated normal resistance to *Aspergillus fumigatus* and *Burkholderia cepacia*, whilst p47^phox-/-^ NADPH oxidase deficient mice were very susceptible to both these organisms [90]. It was suggested that these results cast doubt upon the initial studies describing the importance of these neutral proteases in antimicrobial resistance. However, the mice we constructed and studied were on a 129/SvJ background [84] whereas the background of those studied by Braham Segal and the Shapiro group was C57Black6 [90]. This is important because “Inbred laboratory mouse strains are highly divergent in their immune response patterns as a result of genetic mutations and polymorphisms. Although common inbred mice are considered ‘immune competent’, many have variations in their immune system-some of which have been described-that may affect the phenotype. Recognition of these immune variations among commonly used inbred mouse strains is essential for the accurate interpretation of expected phenotypes, or those that may arise unexpectedly” [91].

### 4.1 Papillon-Lèfevre syndrome (PLS)

This is an interesting and informative syndrome that is characterised by symmetrical palmoplantar hyperkeratosis and periodontal inflammation, causing the loss of both primary and permanent teeth, and pyogenic infection. It is very rare and only about 300-400 cases have been described. It is caused by mutations in the *CTSC* gene, which codes for cathepsin C [92] (also known as dipeptidyl peptidase-1, EC 3.4.14.1) which is a cysteine dipeptidyl aminopeptidase expressed in the cytoplasmic granules of several tissues, with levels highest in lung, macrophages, neutrophils, CD8+ T cells, and mast cells. The enzyme activates cathepsin G, proteinase-3, neutrophil elastase, granzymes A, B, and C, and mast cell chymase and tryptase by removing inhibitory N-terminal dipeptides [93]. It is not clear where and when this dipeptidase acts. It is composed of a dimer of disulfide-linked heavy and light chains, both produced from a single protein precursor, and a residual portion of the propeptide acts as an intramolecular chaperone for the folding and stabilization of the mature enzyme. The quaternary structure of this protein confers this strict dipeptidase activity [94].

Knocking out this gene in mice [95] led to failure of the activation of granzymes A and B, serine proteases most commonly found in the granules of cytotoxic lymphocytes (CTLs), natural killer cells (NK cells) and cytotoxic T cells [96]. In neutrophils this led to an accumulation of neutrophil elastase and a disappearance of cathepsin G with the loss of activity of both. Evidence against the antimicrobial role of neutral proteases has been proffered by the description of a case of the Papillon-Lèfevre syndrome (PLS) whose neutrophils appeared to lack cathepsin G and proteinase 3 but appeared to kill *S. aureus* and *K. pneumoniae* normally [97]. In a similar study of three such cases, concentrations of cathepsin G, neutrophil elastase and proteinase 3 were severely reduced and killing of *E. coli* and *S. aureus* was impaired in two of the three [98] as had been previously described [99]. Killing of *C. albicans* was found to be impaired in all 15 Egyptian cases tested [100]. It is possible that the lack of detection of a killing defect by Sørensen et al. [97] was caused by the method they used to measure bacterial killing. They initially incubated the neutrophils and opsonised bacteria at 37*°*C for 10 minutes, centrifuged them and re-incubated them at 37*°*C for a further 10 and 30 minutes, taking the numbers of bacteria at the start of the second incubation as time 0 and expressing killing as a percentage of the viable bacteria at that time. It is very possible that much of the killing had already occurred in the first 10 minutes and that differences in the function of the patients’ and control cells had been missed.

In addition to the characteristic very aggressive periodontitis, serious infections are common in PLS. 17% of patients have serious cutaneous sepsis and 15 cases of liver abscess, one renal abscess, one case of multiple brain abscesses and a case of pyelonephritis have been described. *S. aureus* and *E. coli* were the main organisms isolated and where documented the histology has been that of a granulomatous inflammation.

A discordant feature is that the periodontitis that characterises PLS is much more severe than in any of the other primary immunodeficiency diseases. It leads to bone resorption around the affected teeth resulting in mobility and loss of most of the teeth. Several organisms have been isolated from the crevicular space of these patients [101], but the bulk of evidence suggests that *Actinobacillus actinomycetemcomitans* species are largely responsible for these infections. Neutrophil serine proteases are important for resistance to these organisms [102] which are killed *in vitro* by neutrophil elastase and cathepsin G [103]. The cathepsin C knock-out mice have no dental problems [104].

It is also of interest that although cathepsin C is thought to be required for the activation of granzymes [95] which are important for T-cell and natural killer cell activity, these patients to not exhibit evidence of T-cell deficiencies such as chronic viral infections.

In addition, in the patient described by Sørensen et al. [97] the neutrophils do produce some cathepsin G and elastase (their Supplemental Table 1), and the lighter, more primitive bone marrow cells produce cathepsin C which disappears in older cells, that in this case had been incubated *in vitro* at 37*°*C for 4 hours before analysis. These young myeloid cells might also contain cathepsin G and elastase, which is important because in serious infections there is a shift to the left of the myeloid series of cells, and these primitive cells enter the circulation [105] and could play an important antimicrobial role.

The interesting feature of this case, which is probably generally applicable to PLS, is why is there a major reduction in the content of these neutral proteases in azurophil granules? The tertiary and quaternary structure of cathepsin C is designed to restrict its activity to that of an exodipeptidase. If this structure were disrupted by activating mutations that lifted this restriction, producing endopeptidase activity, the enzyme, or another protease activated by it, would be able to degrade proteins within the granule, the interior of which has a pH close to the optimum of about 6.0 for cathepsin C [106]. The proteomics data presented by Sørensen et al. [97] indicated that a number of granule proteins, other than the three major neutral proteases, were degraded in their patient’s cells, indicating that digestion of the neutral proteases is not simply due to failure to remove their terminal dipeptide. The release of an active endopeptidase from neutrophils attracted to the periodontal space by infection could digest the periodontal ligament and surrounding bone because the pH is also acidic in this compartment [107].

Based upon the scanty, incomplete and conflicting data from investigations of the very rare Papillon-Lèfevre syndrome patients, the conclusion that “proteases in human neutrophils are dispensable for protection against bacterial infection” [108] appears premature.

## 5 The role of the NADPH oxidase in elevating and regulating the pH of the phagocytic vacuole

Initially the pH of the neutrophil phagocytic vacuole was thought to be acidic [109]. We showed that through the action of the NADPH oxidase the pH actually rises to what was originally thought to be *≈*7.5-8 [82], and these findings have been replicated several times [62], [110]-[112]. In those studies, fluorescein, coupled to phagocytosed organisms, was employed as pH indicator. This is not an ideal indicator for this purpose because it cannot measure changes in pH above *≈*8 [110] and is not generally used in a ratiometric manner. In addition, it has been thought by some to become bleached by the action of MPO in the phagocytic vacuole [54], [113].

In recent studies we have used SNARF [114] to make these measurements [40]. SNARF has a dynamic range of between 6 and 11, is ratiometric, and different preparations can be used to simultaneously measure the pH in the vacuole and cytoplasm. We have shown in several ways that its fluorescent properties are not altered by the conditions in the phagocytic vacuole. In addition, we only measured the pH of *Candida* inside vacuoles, so that the fluorescence signal was not influenced by that of the non-phagocytosed organisms which are maintained at the pH of the buffered extracellular medium. Synchronising the changes in fluorescence to the time of particle uptake, gave a more accurate indication of the temporal pH changes.

Almost immediately following engulfment of the SNARF-labelled *Candida* by human neutrophils the vacuole underwent a significant alkalinisation [40]. The mean maximum pH reached post-phagocytosis was 9.0 (SD 8.3-10.2) and this elevated pH was maintained for 20-30 minutes (Figure 1). When the NADPH oxidase was inhibited by DPI, the vacuole acidified to 6.3 (SD 6.1-6.6) and when DPI was added to cells that had phagocytosed *Candida*, where the vacuolar pH was very alkaline, the vacuolar pH rapidly dropped, which indicates that continued activity of the oxidase is required to maintain this alkaline pH. The addition of 5mM azide had a dramatic effect by acidifying the vacuole by about 2 pH units. This effect was not due to the inhibition of the activity of MPO because it was not observed with two other more specific MPO inhibitors, 4-aminobenzoic acid hydrazide (4ABH) and KCN [40].

**Figure 1:**
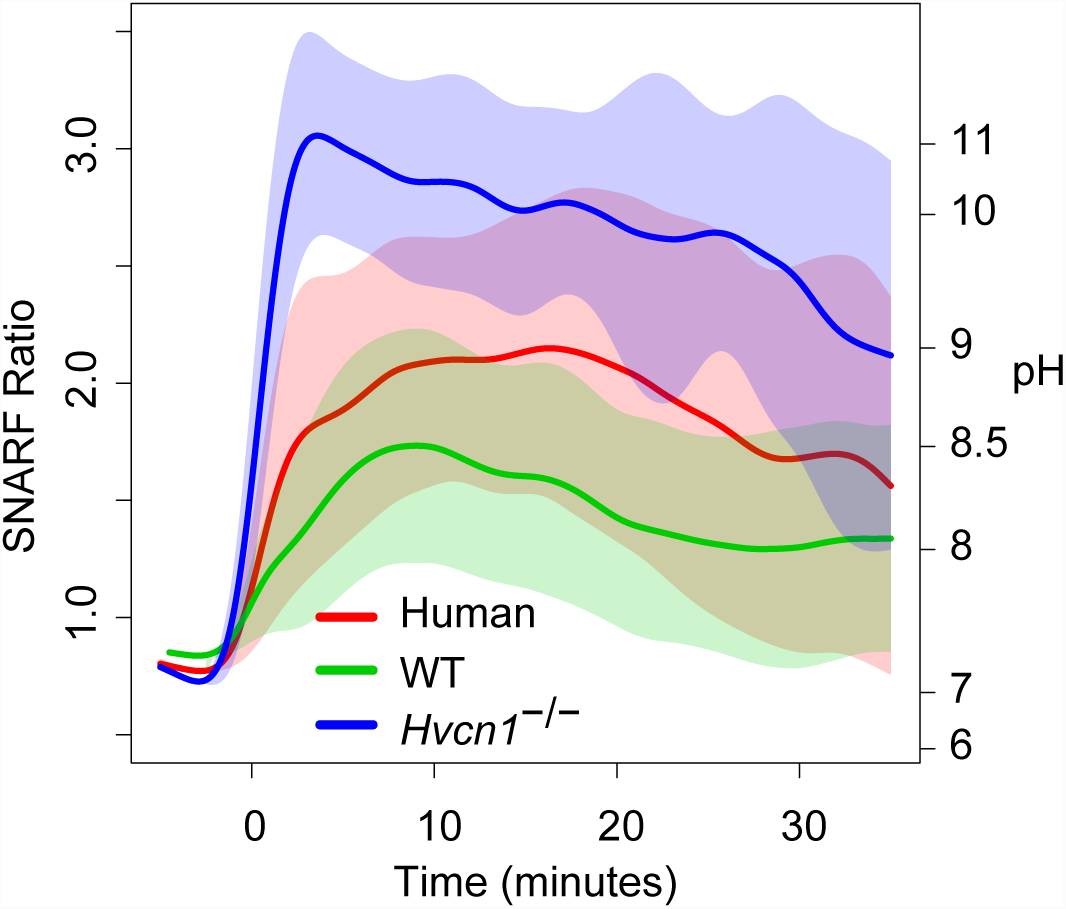
Time courses of changes in vacuolar pH in human, *Hvcn1* ^-/-^ and wild-type mouse neutrophils phagocytosing SNARF labelled *Candida*. From Levine et al. [40]

The observation that the addition of azide to phagocytosing neutrophils produces an acidification of the phagocytic vacuole is important. Previous studies that were unable to detect an elevation of vacuolar pH above neutral in normal neutrophils [115], [116] included azide in all the solutions to counteract bleaching of the fluorescein by MPO. In addition, azide has been included in microbicidal assays to inhibit the action of MPO in order to demonstrate its importance in the killing process [117], [118]. We now know that this inhibition by azide of the killing of microbes by neutrophils could be produced either by blocking the peroxidatic function of MPO, or by impeding the proteolytic activity of the neutral proteases which are much less efficient in the more acidic environment induced by the presence of this agent.

## 6 How is the vacuolar pH elevated?

The transport of electrons into the phagocytic vacuole is electrogenic causing a large, rapid, membrane depolarisation which will itself curtail further electron transport unless there is compensatory ion movement [119] by the passage of cations into the vacuole and/or anions in the opposite direction (Figure 2). The nature of the ions that compensate the charge will have a direct effect on the pH within the vacuole and the cytosol. The cytoplasmic granules are very acidic, with a pH of about 5.5, and they download their acid contents into the vacuole. The electrons that pass into the vacuole produce O_2_^-^ which dismutates to form peroxide (O_2_^2-^) that is then protonated to form H_2_O_2_. The source of these protons will govern the alterations in the vacuolar pH. If all the charge is compensated by protons passing into the vacuole, then none of the protons released into the vacuole from the granules will be consumed and the pH will remain acidic. In fact most of the charge is compensated by protons passing from the cytoplasm into the vacuole through the HVCN1 channel [120], because if this channel is knocked out in mice the vacuolar pH becomes grossly elevated to about 11 (Figure 1) [40]. Under normal physiological conditions about 5-10% of the compensating charge is contributed by non-proton ions, some of which is K^+^ passing into the vacuole [84], the residual ion flux could be the egress of chloride [40]. These non-proton ion channels remain to be identified.

**Figure 2:**
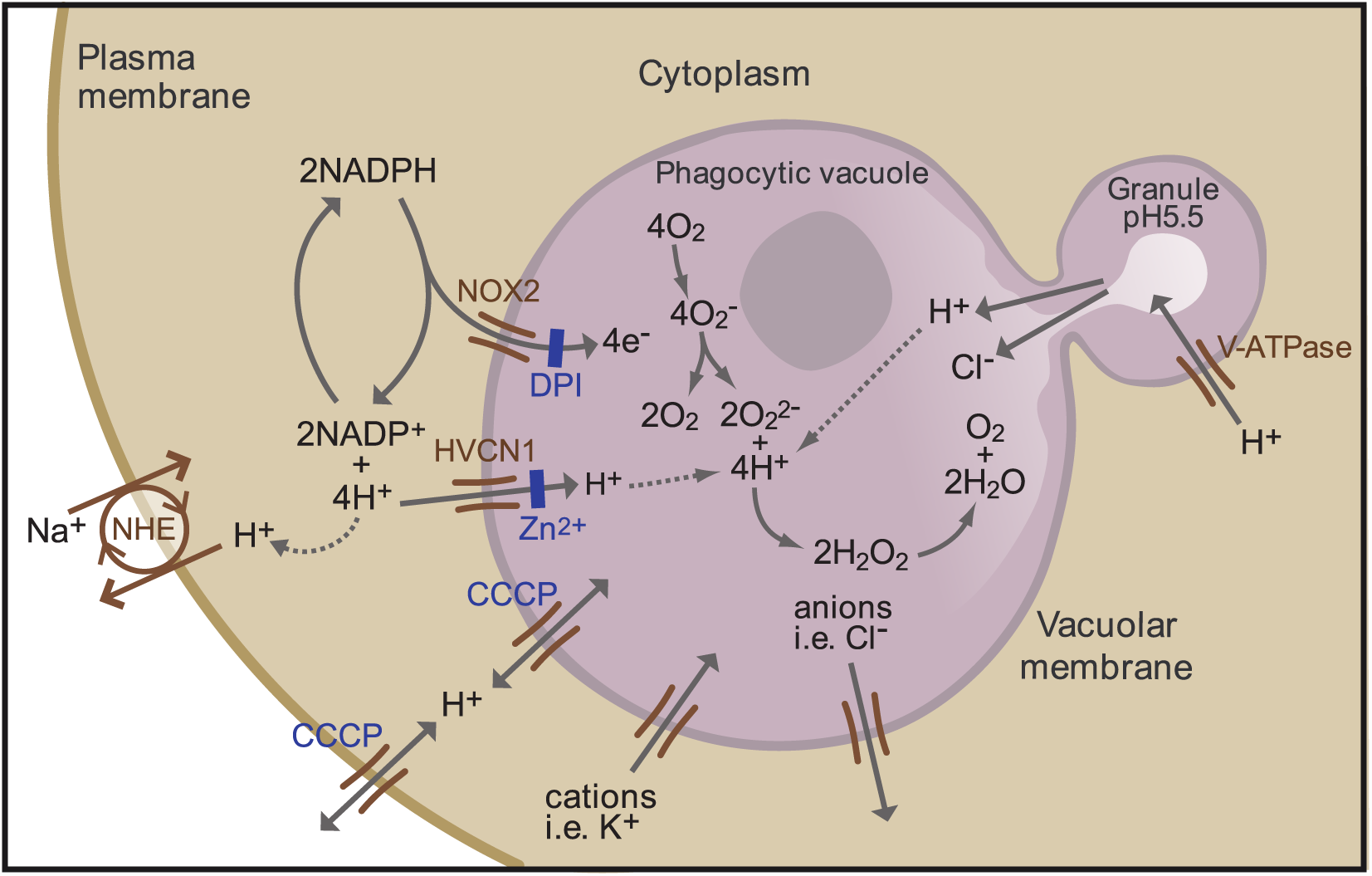
Schematic representation of the neutrophil phagocytic vacuole showing the consequences of electron transport by NOX2 onto oxygen. The proposed ion fluxes that might be required to compensate the movement of charge across the phagocytic membrane together with modulators of ion fluxes are shown. From Levine et al. [40]

## 7 Influence of pH on the activities of cathepsin G, elastase and myeloperoxidase

The accurate measurement of vacuolar pH in normal human neutrophils is absolutely central to understanding the mechanisms by which these cells kill bacteria and fungi. A strong body of opinion supports the concept that MPO is “a front line defender against phagocytosed microorganisms” [53] and that it kills microbes within this compartment through the generation of HOCl. We measured the effect of pH on peroxidase and chlorinating activities of MPO and found these to be were maximal at acidic pH, with both activities falling off substantially as the pH was elevated, until both activities were almost completely abolished at the pH of approximately 9 pertaining in the vacuole [40]. However, as shown here, the elevated pH provides an optimal milieu for the microbicidal and digestive functions of the major granule proteases, elastase, cathepsin G and proteinase 3 (Figure 3), which are activated by this elevated pH and the influx of K^+^ into the vacuole [84].

**Figure 3:**
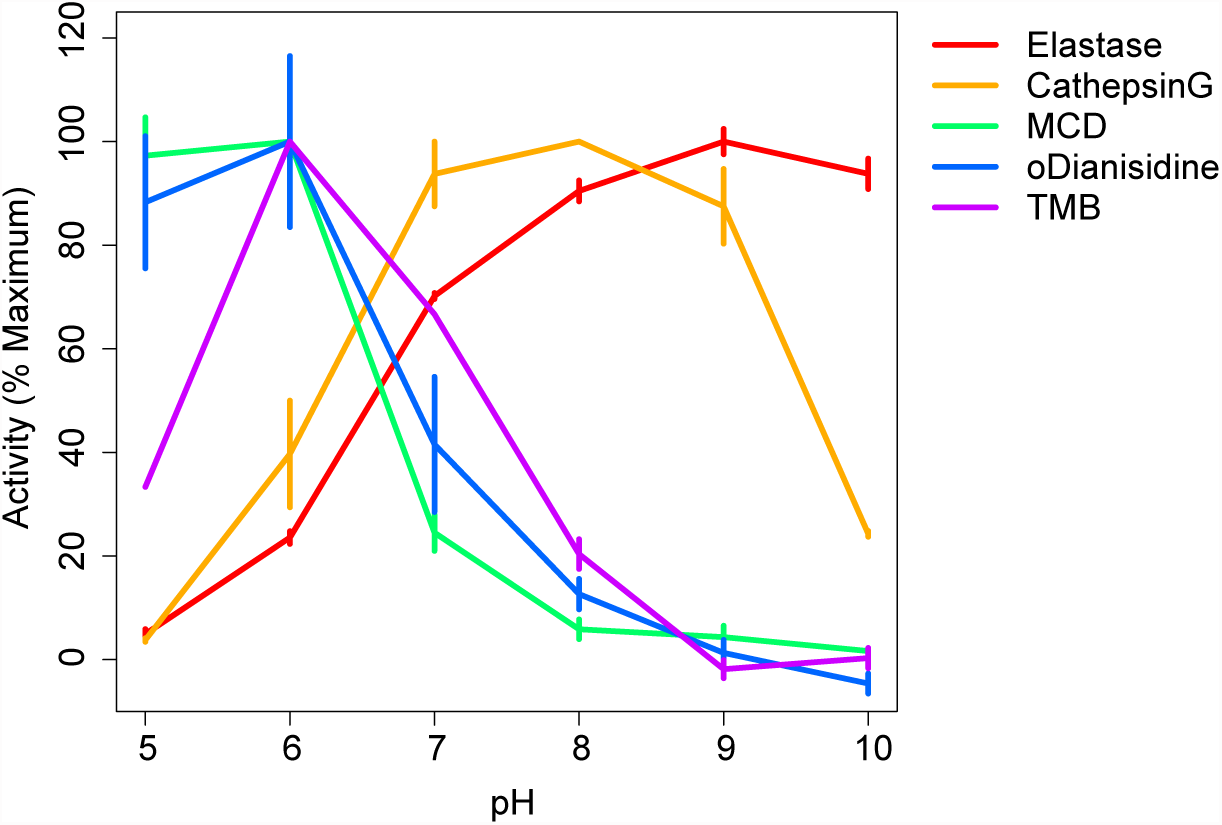
The effect of variations in pH on peroxidatic (TMB and o-Dianisidine) and chlorinating (MCD) activities of MPO and on the protease activities of cathepsin G and of elastase are shown. From Levine et al. [40]

The alkaline vacuolar pH will assist the dissociation of the cationic proteins from the negatively charged sulphated proteoglycan matrix to which they are bound. The pKs of cathepsin G, elastase, proteinase-3 and MPO are all about 10 [121].

In CGD, digestion of ingested microbes is inefficient because the vacuolar pH is very acidic and the granules contents do not disperse within the vacuole [82]. This retained undigested material results in the observed granulomatous tissue response and hyper-inflammatory tissue reactions observed in patients [122] and in experimental animals [123], [124].

These results demonstrate that at the physiological pH pertaining in the phagocytic vacuole, cathepsin G and elastase will be active but that MPO has virtually no activity as a chlorinating peroxidase, although it retains SOD and catalase activities. They also explain the anomalous results produced when azide has been used as an inhibitor of MPO to demonstrate the functional relevance of this enzyme for microbial killing. Azide not only inhibits MPO, but the vacuolar acidification it produces impairs the function of the neutral proteases, so that the effects of azide ascribed to the inhibition of MPO activity might just as well have been, and probably were, due to acidification of the phagocytic vacuole and the negating effect this had on the digestive enzyme activity within this compartment.

It is possible that MPO has evolved to serve two different functions *in vivo*, depending upon the local environment. The pH of *≈*9 in the vacuole favours SOD and catalase [48], [125] activity rather that of a peroxidase, whereas, when the neutrophil encounters an organism that it is unable to fully engulf, such as a fungal mycelium, the situation is different [126]. The concentration of MPO and H_2_O_2_ will be much lower, the pH, which is low in inflammatory foci at *≈*6 [75] is optimal for peroxidatic activity, and the supply of chloride is limitless.

## 8 The family of NOXs and DUOXs

With the development of high-throughput DNA sequencing, and efficient computer based analysis and comparison of sequences, it emerged that a large family of NADPH oxidases, termed NOXs, exist throughout the plant and animal kingdoms [127]-[129]. All these NOXs conformed to the structural organisation of the prototype gp91^phox^ [25], [27], [130], which inexplicably was called NOX2 [127] even though it was discovered two decades before the other NOXs. Variants of the NOXs are the DUOXs which have a very similar structure but have additional domains with homology to peroxidases, but which act as superoxide dismutases, causing the DUOXs to generate H_2_O_2_ rather than O_2_^-^ [131].

### 8.1 NOX3 and inner ear function

The NOX3 knock-out mouse gives important clues as to the general function of NOXs. These mice lack normal spatial awareness because of abnormalities of their otoconia [132]. These small bodies are structures in the saccule and utricle of the inner ear, and give information as to position and changes in movement. Mass is added to these proteinaceous bodies by the deposition of calcium carbonate, and it is the failure to calcify their otoconia that produces the malfunction in the NOX3 mouse. The question is why this failure of otoconial calcification occurs. The ionic composition of the endolymph of the inner ear has similarities with that of the vacuole, having a high concentration of K^+^ and an alkaline pH [133], [134], the latter being required for the deposition of CaCO_3_, the solubility of which is very pH dependent. It is probable that NOX3 plays an important part in the establishment or maintenance of these ionic conditions.

### 8.2 NOXs in plants

The NOXs in plants are called RBOHs (Respiratory Burst Oxidase Homologues) and are important for the plant defence response [135]. They also have more specific functions in specialised organelles of the plant, including the pollen tubes, lateral roots and stomatal guard cells as examples. These cells all contain large vacuoles surrounded by membranes containing a variety of ion channels. In general, K^+^ enters these vacuoles increasing the osmotic pressure and causing growth of the pollen tube [136], [137], extension of lateral roots [138] and stomatal opening [139].

### 8.3 NOXs in fungi

There is growing evidence that NOXs are important for many aspects of fungal life including vegetative hyphal growth, differentiation of conidial anastomosis tubes, fruiting body and infection structure formation, and for the induction of apoptosis [140].

## 9 Conclusion

With the discovery of the production of oxygen free radicals by neutrophils, free radical chemistry was transported from the egis of the radiation chemist to that of the cell biologist and clinical investigator. The initial concept was that these oxygen radicals were very toxic, and that they themselves could be held responsible for microbial killing in professional phagocytes, and for tissue damage at sites of inflammation.

After a lot of work, and a clearer understanding of free radical reactivity and toxicity, together with clinical trials of radical scavengers and antioxidants that proved therapeutically ineffective [141] [142], a consensus has emerged that these molecules are not as toxic as was originally envisaged. In the case of the neutrophil the mantle of a toxic mechanism reverted to that of MPO mediated halogenation. However, it was subsequently discovered that these NOXs are widely distributed throughout the biological world, and a function, or functions had to be ascribed to them. With the observation that the production of these reduced oxygen products (ROS) could be induced by a stimulus, and that another cellular response was also evoked, and in the absence of NOX activity this additional response was abrogated, several functions, including that of signalling molecules has been given to these ROS.

Careful investigation has demonstrated that the neutrophil oxidase has a dramatic effect upon the pH and ionic composition of the phagocytic vacuole which in turn influences the functional efficiency of the myriad of enzymes released into this compartment from the cytoplasmic granules. This has a direct effect of the efficiency with which these enzymes kill and digest the phagocytosed microbes.

With the phagocytic vacuole as an example, the function of the NOXs housed in different biological niches can be examined in a different light. Rather than simply assuming that knock-out or knock-down of NOXs at different sites in plants exert their effect by the removal of a signalling mechanism, examination of the effect of the removal of an electromotive force that produces ion fluxes with secondary osmotic and concomitant mechanical forces needs to be considered, together with the consequent pH changes.

In summary, the NOXs provide a simple and highly efficient electrochemical generator. The pathways to the generation of the substrate, NADPH, are metabolically efficient, particularly in plants where it is a primary product of photosynthesis. The NOX enzymes contain one molecule of FAD and two haems, one close to the inner and the other to the outer surface of the membrane, and the acceptor is oxygen, which is largely ubiquitous. This simple system generates an electromotive force that can be used to drive cations in the same direction as the electrons, or anions in the opposite direction. The separation of electrons and protons alters the pH on both sides of the membrane. These physicochemical changes can be adapted to a variety of biological applications.

## Acknowledgements

We thank the Wellcome Trust, Medical Research Council, Charles Wolfson Charitable Trust and Irwin Joffe Memorial Trust for financial support

